# Interplay between Skin Sympathetic Nerve Activity and Gut Microbiota, the Neurocardiac Axis in Acute Coronary Syndrome

**DOI:** 10.1101/2025.05.06.652555

**Authors:** Wei-Chung Tsai, Po Peng, Pei-Syuan Jhou, Yi-Chun Tsai, Wei-Wen Hung, Yi-Hsiung Lin, Shih-Jie Jhuo, Tien-Chi Huang, Li-Fang Chou, Tsung-Hsien Lin, Ho-Ming Su, Wen-Ter Lai, Chien-Hung Lee, Bin-Nan Wu, Shien-Fong Lin, Wei-Chun Hung

## Abstract

**BACKGROUND:** Previous studies have linked gut dysbiosis to the acute coronary syndrome (ACS) and sympathetic nervous system disturbances. However, the relationship between skin sympathetic nerve activity (SKNA) and gut microbiota in ACS patients remains largely unexplored. We hypothesize that elevated SKNA may interact with specific microbial profiles in ACS.

**METHODS AND RESULTS:** Fifteen ACS patients with age- and sex-matched controls were enrolled in the cross-sectional study. Subjects with type 2 diabetes or on antibiotics, proton pump inhibitors, or metformin were excluded. Demographic and clinical data were recorded, and SKNA was measured by neuECG. Gut microbial profiles were analyzed by sequencing the V3-V4 region of the 16S rRNA gene. Significant differences in alpha diversity, including Chao 1 index and Faith’s phylogenetic diversity, were observed between the ACS and control groups (*P*<0.001). Distinct beta diversity between the two groups was also identified using unweighted normalized UniFrac analysis (*P*=0.001). Redundancy analysis and Mantel test revealed a significant relationship between SKNA and microbial profiles. Several butyrate-producing bacteria were decreased in the ACS group, while the genus *Hungatella*, a trimethylamine producer, showed a significant increase and was positively correlated with SKNA. Additionally, predicted microbial functional analysis indicated an elevation in branched-chain amino acid biosynthesis in the ACS group.

**CONCLUSIONS:** This study is the first to establish an association between SKNA and gut microbial profiles in ACS patients. SKNA can serve as a biomarker for ACS, offering a simple and non-invasive tool to explore the interaction between the nervous system and gut microbiota in cardiovascular diseases.

**Clinical Perspective:** *WHAT IS NEW?:* The Skin sympathetic nerve activity (SKNA) is a novel and non-invasive recording of the sympathetic nervous system from the skin of patients. The NEW findings in our study from acute coronary syndrome (ACS) patients are (1) to establish the correlation between SKNA, heart rate variability and microbiota, (2) the *Hungatella* is correlated with SKNA and standard deviation of NN intervals (SDNN).

*WHAT ARE THE CLINICAL IMPLICATIONS?:* Since microbiota and sympathetic nervous system (SNS) might be essential factors of ACS, it is important to assess the abundance of microbiota and SNS activity for accurate management of ACS. The *Hungatella*, one of the microbiota involved in atherosclerotic cardiovascular disease, heart failure, inflammatory process, and anxiety disorder, might be the link between SNS and ACS. In our study, we prove the positive association between *Hungatella* and SKNA. Therefore, by checking the SKNA of patients with ACS, we might discover the SNS activity and also the underlying microbiota etiology of ACS. Then, it is critical and accurate to target *Hungatella* in ACS patients with high SKNA to reduce the risk of ACS.

## Introduction

Acute coronary syndrome (ACS) is one of the leading causes of mortality worldwide.^1^ Hemodynamically significant ventricular arrhythmias (VA) during ACS occur in approximately 8% of patients and lead to mortality.^2^ The sympathetic hyperinnervation caused by ACS can lead to the development of VA. Sympathetic drive leads to increased automaticity, triggered activity, and re-entry mechanisms in the heart, all of which can contribute to VA.^3,4^ Sympathetic overdrive after ACS could provoke further myocardial ischemia, leading to a vicious cycle of VA and sudden cardiac death. Heart rate variability (HRV), heart rate turbulence, and abnormal T wave alternans can be used to assess the risk of death or VA after MI.^5,6^ neuECG is a well-validated non-invasive method of simultaneously recording electrocardiogram and skin sympathetic nerve activity (SKNA).^7^ Increased SKNA is known to precede spontaneous VA in canine models and humans.^8–11^ We also found that SKNA increased in ACS patients and correlated with VA in a clinical study.^12^

Accumulating evidence has demonstrated the critical roles of gut dysbiosis and the imbalance of their metabolites in the development of ACS. A decrease in butyrate-producing bacteria, along with a corresponding reduction in butyrate concentrations, is correlated with elevated inflammation, cardiac damage, and increased mortality.^13^ Trimethylamine N-oxide (TMAO), which is formed from trimethylamine (TMA) generated by the action of gut microbiota, has been reported as a novel prognostic marker for ACS.^14^ Also, the relationships between gut microbiota and circulating metabolites demonstrated by Liu et al. provide a better understanding of the interplay between host and gut microbiota in atherosclerotic pathogenesis.^15^ A canine study shows that TMAO can activate the cardiac sympathetic nervous system (SNS) and worsen ischemic VA via the left stellate ganglion (SG) pathway or central autonomic activation.^16^ Accumulation of branched-chain amino acids (BCAA) is an alternative biomarker of cardiometabolic disease.^17–19^ Pedersen et al. further suggest that elevated serum BCAA levels were attributed to gut microbiome rather than dietary intake.^20^

As mentioned above, gut microbiota have been reported to play a role in ACS and could potentially modulate sympathetic neurons. However, the interplay between the neuro-cardiac axis and the specific microtome has not been clearly understood in clinical ACS patients. Considering all the above evidence, we conducted the study to test the hypothesis that SKNA is increased in ACS patients compared to controls and that the increased SKNA is associated with specific microtomes that may play an important role in linking microbiota, SNS, and ACS.

## METHODS

### Participants

This single-center, cross-sectional study was approved by the ethics committee of Kaohsiung Medical University Hospital, and all study subjects provided written informed consent. This study is registered with ClinicalTrials.gov, identifier: NCT03243448. The enrollment period of the study participants was from 2019 to 2021 at Kaohsiung Medical University and Kaohsiung Medical University Hospital. Inclusion criteria: ACS patients and control participants who were 1:1 matched in terms of age and gender. Each group consisted of 15 individuals, all 20 years or older. ACS patients were defined as those with myocardial infarction and unstable angina admitted to the coronary care unit after coronary angiography and/or percutaneous coronary intervention as standard of care. The controls enrolled were healthy adults who joined the health management program at Kaohsiung Medical University. Exclusion criteria for this study included the use of antibiotics, metformin, or proton pump inhibitors, and diagnoses of diabetes or colon cancer, as all these could interfere with the results of the microbiota study. Indeed, the stringent exclusion criteria can make enrolling ACS patients in the study challenging. Demographic and clinical data such as age, gender, body mass index (BMI), clinical factors (hypertension, diabetes, dyslipidemia, mean arterial pressure) and laboratory data (estimated glomerular filtration rate, serum level of potassium, triglyceride, low-density lipoprotein (LDL) cholesterol, high-density lipoprotein (HDL) cholesterol, hemoglobin, hemoglobin A1C were collected in both ACS and control groups.

### neuECG, SKNA, and HRV Measurements

The detailed methods of neuECG recording have been reported elsewhere.^7^ In brief, the neuECG used conventional lead I ECG electrodes and equipment with a high sampling rate (10,000 Hz) and wide bandwidth (1-2000 Hz) version of the ME6000 Biomonitor System (Mega Electronics Ltd, Finland) to record the electrical signal from the skin of the research subjects. The signals were then band-pass filtered at 500-1000 Hz to display SKNA and 1-150 Hz to display ECG. The neuECG was recorded during baseline, stress^21^, and recovery (5 minutes for each phase). The stress consisted of mental arithmetic stress induced by arithmetic involving the subtraction of serial 13’s from a 4-digit number for 5 minutes. Data were analyzed using customized software to determine the average SKNA (aSKNA [µV]) per digitized sample over the monitoring period.^7^ The research subjects were asked to lie down and rest for 10 minutes prior to SKNA recording to reduce bias due to motion artifacts and SNS interference..

Short-term HRV analysis was conducted for 5-minute intervals during each phase of SKNA recording, in order to facilitate comparison. The R peak of the QRS complex in each 5-minute window of the neuECG signal was automatically detected by the modified Pan Tompkins algorithm^22^ and the RR interval was obtained beat by beat. We analyzed the time and frequency domain of HRV using MATLAB (Mathworks, Inc., USA).^22,23^

### Fecal Sample Collections

The fecal samples were frozen immediately after collection and delivered to the laboratory in a cooler bag within 24 hours. Subsequently, the samples were stored at -80 °C for up to 3 days before processing.

### Fecal DNA Extraction and 16S rRNA Sequencing

Fecal DNA was extracted using a stool DNA extraction kit (Topgen Biotechnology Co., Ltd., Kaohsiung, Taiwan) as previously described.^24^ The DNA concentration and quality were assessed using the Colibri Microvolume spectrophotometer (Titertek Berthold, Pforzheim, Germany) before frozen at −20 °C to preserve its integrity.

The V3-V4 regions of 16S rRNA gene in each sample were amplified using the primer pairs 341F (5’- TCGTCGGCAGCGTCAGATGTGTATAAGAGACAGCCTACGGGNGGCWGCAG -3’) and 805R (5’- GTCTCGTGGGCTCGGAGATGTGTATAAGAGACAGGACTACHVGGGTATCTA ATCC -3’). Library preparation was conducted using the Illumina MiSeq platform, generating 2 x 300 bp reads. Raw sequence data was imported into QIIME2,^25^ where paired-end reads were merged and denoised into amplicon sequence variants (ASVs) using the DADA2 plugin,^26^ as previously described.^24^

### Statistical Analysis

Disease status (ACS vs. health control) was our primary comparison variable, with the baseline SKNA value as a continuous outcome. A priori sample size calculation, assuming a two-sided type I error of 5%, a standard deviation of 0.25 for SKNA, and a 1:1 case/control ratio, estimated a total of 28 participants (14 ACS and 14 controls) to achieve 80% statistical power for detecting an effect size of 0.28.

The significance of differences in anthropometric data, clinical factors, laboratory results, SKNA parameters, and HRV parameters between the ACS and control groups was assessed using SPSS version 18.0 for Windows (SPSS Inc., Chicago, Illinois). The *P* values were calculated using the Mann-Whitney *U* test for continuous data and the Chi-square test for categorical data. A *P* value of <0.05 was considered statistically significant.

### Bioinformatics

We employed the Chao 1 index and Faith’s phylogenetic diversity to assess alpha diversity, comparing differences between the ACS and control groups using Kruskal-Wallis tests. For beta diversity, we performed analysis of similarities (ANOSIM) and permutational multivariate analysis of variance (PERMANOVA) with 999 permutations. These analyses were assessed using principal coordinate analyses (PCoA) based on unweighted UniFrac measurements.^27^ Additionally, redundancy analysis was conducted to examine the relationship between microbial beta diversity and clinically relevant factors.

ASV taxonomy assignment was conducted using SciKit Learn-based approach,^28^ querying the SILVA reference database (release v138, trimmed to the V3-V4 region, L7 taxonomy).^29^ Significant different taxa between the ACS and control groups were identified using linear discriminant analysis effect size (LEfSe) with *P* < 0.05 (factorial Kruskal-Wallis test).^30^ To enhance the stringency, the logarithmic linear discriminant analysis (LDA) score was set to 3. Moreover, random forest analysis was employed to identify the discriminant genera between the two groups.

The fixed-effects models explored the relationships between the targeted genera and SKNA and the standard deviation of NN intervals (SDNN). Generalized linear mixed-effects models (GLMMs) with sequencing depth as weights were fitted using the lme4 package in R. Age and gender were included as random effects.

Phylogenetic investigation of communities by reconstruction of observed states 2 (PICRUSt2) was utilized to predict functional differences between the microbial communities of the ACS and control groups, based on the MetaCyc database.^31^

## RESULTS

### Clinical Characteristics

A total of 15 patients with ACS were included in this study, along with the matched control subjects. The clinical characteristics are shown in Table 1. The two groups had no statistically significant differences in anthropometric data and clinical factors. For the laboratory data, the ACS group exhibited significantly elevated fasting glucose and HbA1c levels and markedly reduced HDL levels compared to the control group. The SKNA parameters during baseline and recovery phases (SKNAb and SKNAr) were significantly higher in the ACS group. Additionally, the ACS group demonstrated significantly diminished SDNN values during the stress phase (SDNNs) in the HRV parameters.

**Table 1.**
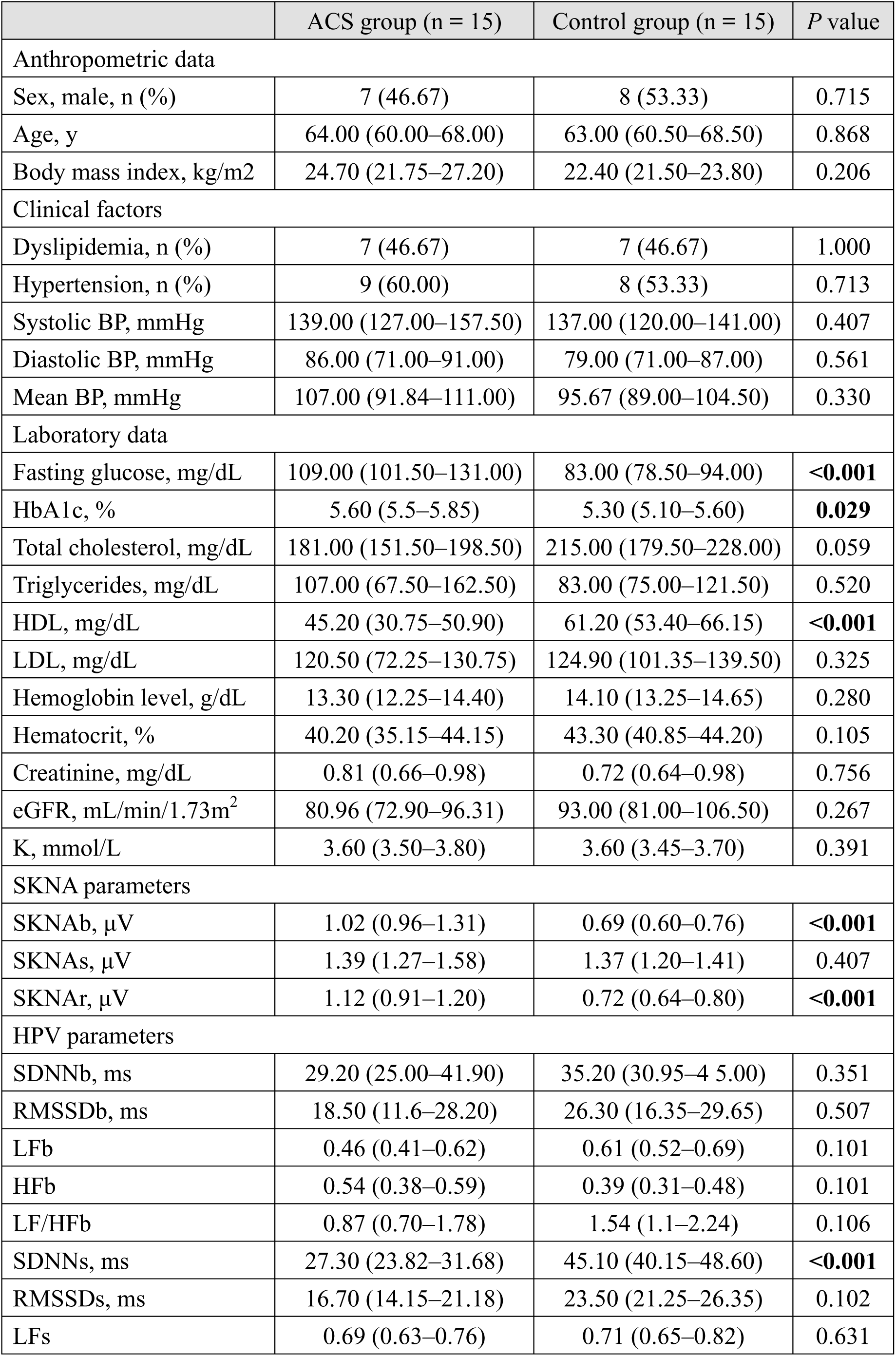
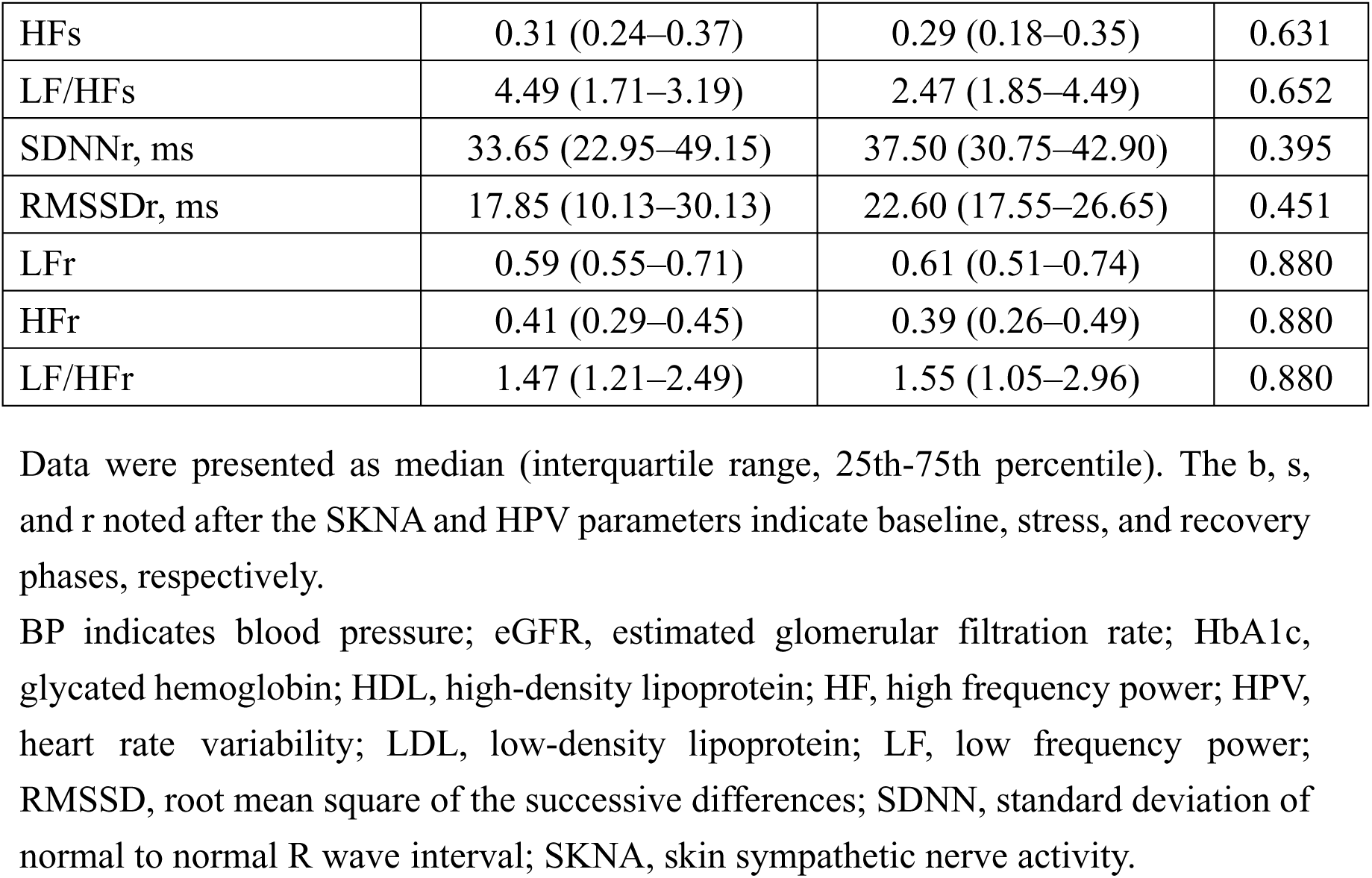
Characteristics of the ACS and control groups.

|  | ACS group (n = 15) | Control group (n = 15) | *P* value |
| --- | --- | --- | --- |
| Anthropometric data |  |  |  |
| Sex, male, n (%) | 7 (46.67) | 8 (53.33) | 0.715 |
| Age, y | 64.00 (60.00–68.00) | 63.00 (60.50–68.50) | 0.868 |
| Body mass index, kg/m<sup>2</sup> | 24.70 (21.75–27.20) | 22.40 (21.50–23.80) | 0.206 |
| Clinical factors |  |  |  |
| Dyslipidemia, n (%) | 7 (46.67) | 7 (46.67) | 1.000 |
| Hypertension, n (%) | 9 (60.00) | 8 (53.33) | 0.713 |
| Systolic BP, mmHg | 139.00 (127.00–157.50) | 137.00 (120.00–141.00) | 0.407 |
| Diastolic BP, mmHg | 86.00 (71.00–91.00) | 79.00 (71.00–87.00) | 0.561 |
| Mean BP, mmHg | 107.00 (91.84–111.00) | 95.67 (89.00–104.50) | 0.330 |
| Laboratory data |  |  |  |
| Fasting glucose, mg/dL | 109.00 (101.50–131.00) | 83.00 (78.50–94.00) | **<0.001** |
| HbA1c, % | 5.60 (5.5–5.85) | 5.30 (5.10–5.60) | **0.029** |
| Total cholesterol, mg/dL | 181.00 (151.50–198.50) | 215.00 (179.50–228.00) | 0.059 |
| Triglycerides, mg/dL | 107.00 (67.50–162.50) | 83.00 (75.00–121.50) | 0.520 |
| HDL, mg/dL | 45.20 (30.75–50.90) | 61.20 (53.40–66.15) | **<0.001** |
| LDL, mg/dL | 120.50 (72.25–130.75) | 124.90 (101.35–139.50) | 0.325 |
| Hemoglobin level, g/dL | 13.30 (12.25–14.40) | 14.10 (13.25–14.65) | 0.280 |
| Hematocrit, % | 40.20 (35.15–44.15) | 43.30 (40.85–44.20) | 0.105 |
| Creatinine, mg/dL | 0.81 (0.66–0.98) | 0.72 (0.64–0.98) | 0.756 |
| eGFR, mL/min/1.73m<sup>2</sup> | 80.96 (72.90–96.31) | 93.00 (81.00–106.50) | 0.267 |
| K, mmol/L | 3.60 (3.50–3.80) | 3.60 (3.45–3.70) | 0.391 |
| SKNA parameters |  |  |  |
| SKNAb, μV | 1.02 (0.96–1.31) | 0.69 (0.60–0.76) | **<0.001** |
| SKNAs, μV | 1.39 (1.27–1.58) | 1.37 (1.20–1.41) | 0.407 |
| SKNAr, μV | 1.12 (0.91–1.20) | 0.72 (0.64–0.80) | **<0.001** |
| HPV parameters |  |  |  |
| SDNNb, ms | 29.20 (25.00–41.90) | 35.20 (30.95–45.00) | 0.351 |
| RMSSDb, ms | 18.50 (11.6–28.20) | 26.30 (16.35–29.65) | 0.507 |
| LFb | 0.46 (0.41–0.62) | 0.61 (0.52–0.69) | 0.101 |
| HFb | 0.54 (0.38–0.59) | 0.39 (0.31–0.48) | 0.101 |
| LF/HFb | 0.87 (0.70–1.78) | 1.54 (1.1–2.24) | 0.106 |
| SDNNs, ms | 27.30 (23.82–31.68) | 45.10 (40.15–48.60) | **<0.001** |
| RMSSDs, ms | 16.70 (14.15–21.18) | 23.50 (21.25–26.35) | 0.102 |
| LFs | 0.69 (0.63–0.76) | 0.71 (0.65–0.82) | 0.631 |
| HF<sub>s</sub> | 0.31 (0.24–0.37) | 0.29 (0.18–0.35) | 0.631 |
| LF/HF<sub>s</sub> | 4.49 (1.71–3.19) | 2.47 (1.85–4.49) | 0.652 |
| SDNN<sub>r</sub>, ms | 33.65 (22.95–49.15) | 37.50 (30.75–42.90) | 0.395 |
| RMSSD<sub>r</sub>, ms | 17.85 (10.13–30.13) | 22.60 (17.55–26.65) | 0.451 |
| LF<sub>r</sub> | 0.59 (0.55–0.71) | 0.61 (0.51–0.74) | 0.880 |
| HF<sub>r</sub> | 0.41 (0.29–0.45) | 0.39 (0.26–0.49) | 0.880 |
| LF/HF<sub>r</sub> | 1.47 (1.21–2.49) | 1.55 (1.05–2.96) | 0.880 |

Data were presented as median (interquartile range, 25th-75th percentile). The b, s, and r noted after the SKNA and HPV parameters indicate baseline, stress, and recovery phases, respectively.
BP indicates blood pressure; eGFR, estimated glomerular filtration rate; HbA1c, glycated hemoglobin; HDL, high-density lipoprotein; HF, high frequency power; HPV, heart rate variability; LDL, low-density lipoprotein; LF, low frequency power; RMSSD, root mean square of the successive differences; SDNN, standard deviation of normal to normal R wave interval; SKNA, skin sympathetic nerve activity.

### The ACS Patients Displayed Altered Gut Microbiota Composition Compared to Healthy Controls

The composition of gut microbiota in the ACS and control groups was determined using the 16S rRNA gene sequencing. As depicted in Figure 1, a notable reduction in alpha diversity was observed in the ACS group, evident from the Chao 1 index (*P* value < 0.001) and Faith’s phylogenetic diversity (*P* value < 0.001). Distinct beta diversity clustering of gut microbiota between the ACS and control groups was observed using unweighted UniFrac (ANOSIM: R =0.702, *P* value = 0.001; PERMANOVA: pseudo-F = 9.573, *P* value = 0.001) or weighted unnormalized UniFrac measurements (ANOSIM: R = 0.123, *P* value = 0.008; PERMANOVA: pseudo-F = 2.860, *P* value = 0.011), as illustrated in Figure 2.

**Figure 1.**
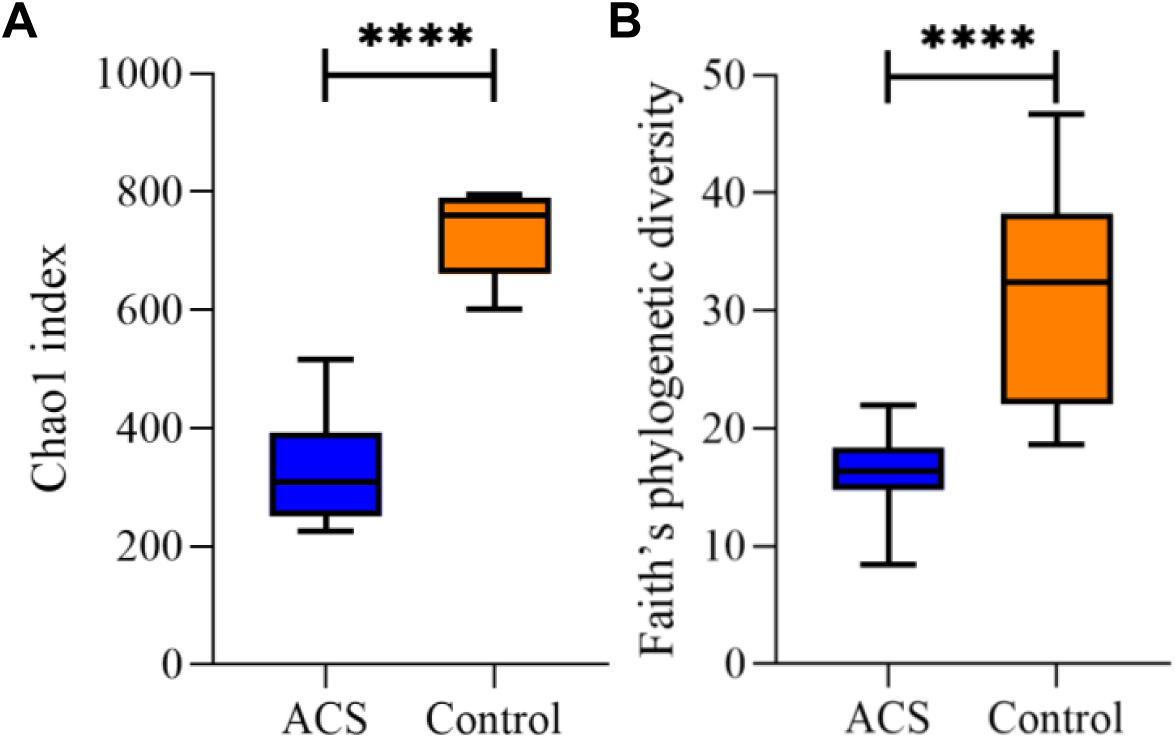
Comparison of the alpha diversity between the ACS and control groups. **A**, the median of Chao1 index was 309.00 (25th-75th percentile: 257.15-383.27) in the ACS group and 760.38 (671.15-783.42) in the control group. **B**, the median of Faith’s phylogenetic diversity was 16.39 (25th-75th percentile: 14.81-18.20) in the ACS group and 32.46 (23.73-37.83) in the control group.

**Figure 2.**
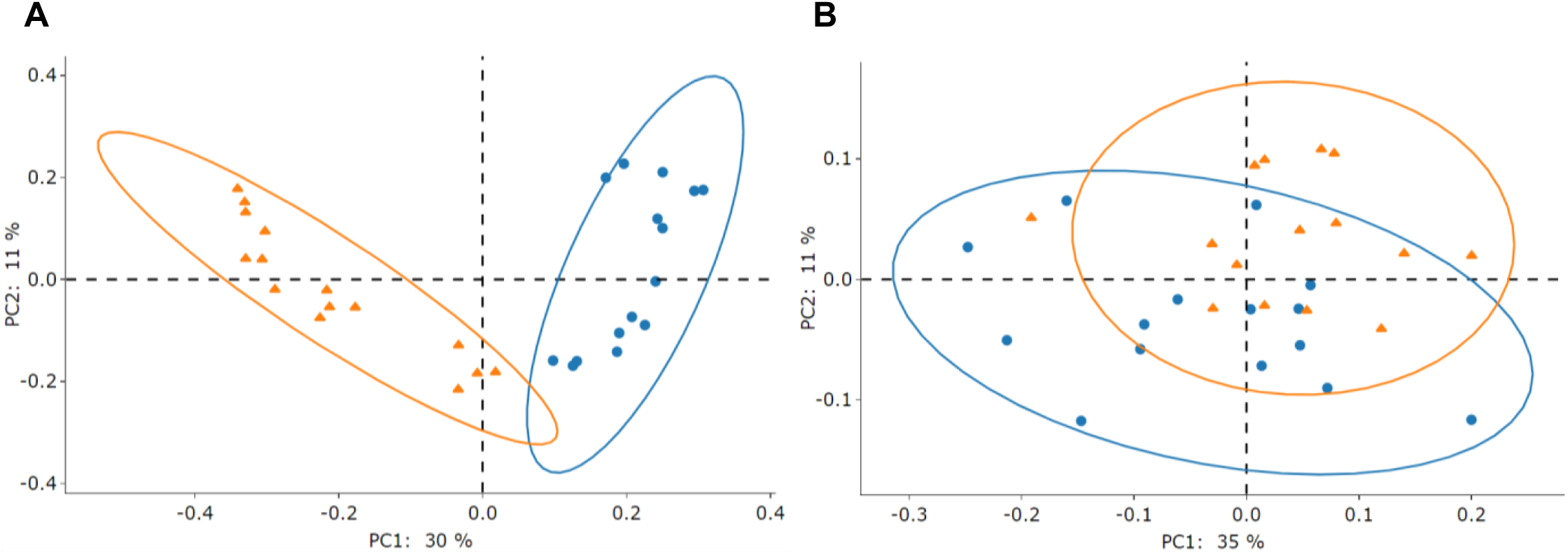
Beta diversity clustering of gut microbiota between the ACS and control groups. **A**, the unweighted UniFrac measurements. **B**, the weighted unnormalized UniFrac measurements. Blue circle indicates the ACS group, and orange triangle indicates the control group.

### Association between clinically relevant variables and microbial beta diversity

Clinically relevant variables were further investigated for their association with microbial beta diversity by RDA. Fasting glucose was excluded from further analysis due to stress hyperglycemia in the ACS subjects despite significant differences between the ACS and control groups. To mitigate the potential accumulation of signals, the SKNA data were segregated into three distinct phases: baseline, stress, and recovery (Figures 3A to 3C). The SKNAb, SKNAr, SDNNs, and HDL had the most potent influences (the longest arrow) on microbial beta diversity. The SKNAb and SKNAr showed positive correlations with the ACS group during the baseline and recovery phases, respectively. Conversely, SDNNs negatively correlated with the ACS group in the stress phase. The HDL levels positively correlated with the control group in all three phases.

**Figure 3.**
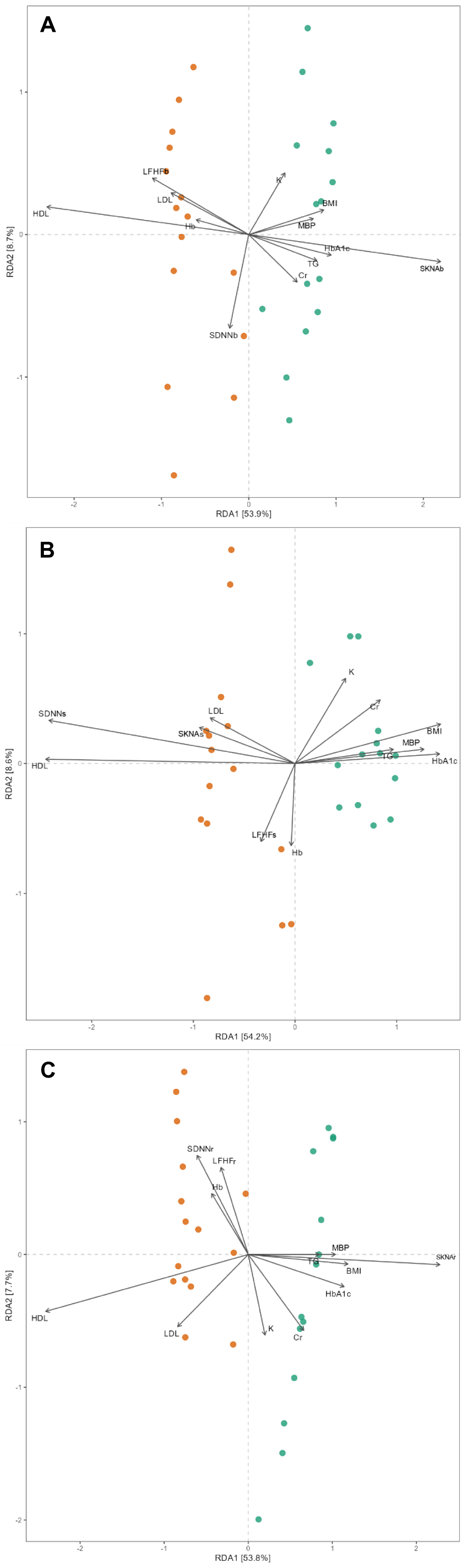
Clinical relevant factors influencing the beta microbial beta diversity between the ACS and control groups. The clinical confounding factors of BMI (body mass index), Cr (creatinine), Hb (hemoglobin), HbA1c, HDL (high-density lipoprotein), K (potassium), LDL (low-density lipoprotein), LF/HF (low frequency power/high frequency power), MBP (mean blood pressure), SDNN (standard deviation of normal to normal R wave) and SKNA (skin sympathetic nerve activity) were examined by redundancy analysis (RDA) based on the unweighted UniFrac measurements. **A**, the baseline phase. **B**, the stress phase. **C**, the recovery phase. Green spot indicates the subjects belonging to the ACS group, and orange spot indicates the subjects belonging to the control group.

Mantel test further confirmed that the SKNAb (*r* = 0.151, *P* value = 0.023), SDNNs (*r* = 0.249, *P* value = 0.002), and SKNAr (*r* = 0.209, *P* value = 0.007) were also significantly related to gut microbiota profiles during the baseline, stress, and recovery phases, respectively (Table 2). The HDL levels were significantly associated with the dissimilarity of gut microbiota in all three phases. No other clinical parameters showed a significant relationship with microbial beta diversity.

**Table 2.** Evaluation of the relationship between clinical relevant factors and gut microbial profiles by Mantel test. HDL indicates high-density lipoprotein; SDNN, standard deviation of normal to normal R wave interval; SKNA, skin sympathetic nerve activity.

|  | Phase |  |  |  |  |  |
| --- | --- | --- | --- | --- | --- | --- |
|  | Baseline |  | Stress |  | Recovery |  |
|  | Correlation coefficient | *P* value | Correlation coefficient | *P* value | Correlation coefficient | *P* value |
| SKNA | 0.151 | **0.023** | -0.014 | 0.522 | 0.209 | **0.007** |
| SDNN | -0.028 | 0.603 | 0.249 | **0.002** | -0.090 | 0.942 |
| HDL | 0.273 | **0.003** | 0.235 | **0.002** | 0.262 | **0.002** |

### Relative abundances of fecal microbiota between the ACS and control groups

Relative differences in the abundances of fecal microbiota between the ACS and control groups were estimated by LEfSe, employing a logarithmic LDA score threshold of > 3. Within the ACS group was an enrichment of the phylum Desulfobacterota, the class Desulfovibrionia, as well as 3 orders, 6 families, 15 genera, and 19 species (Figure S1). Within the control group, enrichment was observed in the phylum Bacteroidota, the classes Actinobacteria and Bacteroidia, as well as 3 orders, 7 families, 29 genera, and 45 species. Remarkably, *Bifidobacterium bifidum*, classified as a lactic acid bacterium, exhibited enrichment extending throughout its taxonomic hierarchy up to the class level in the control group.

Random forest analysis was employed to enhance the stringency of identifying discriminant taxa, and 22 genera were found to be discriminant by both the LEfSe and random forest analysis (Figure 4A). *Eisenbergiella*, *Hungatella*, *Ligilactobacillus*, and Ruminococcaceae UBA1819 were enriched in the ACS group. At the same time, 18 genera were decreased, including four genera belonging to Prevotellaceae (*Paraprevotella*, *Prevotella*, *Prevotella*_9, and Prevotellaceae_NK3B31_group) and several butyrate producers (including *Anaerostipes*, *Butyricicoccus*, *Catenibacterium*, *Eubacterium ruminantiu*m_group, *Fusicatenibacter*, *Lachnospira*, and *Roseburia*). Additionally, the ACS group observed an unexpected decrease in opportunistic microbes *Haemophilus* and *Klebsiella*.

**Figure 4.**
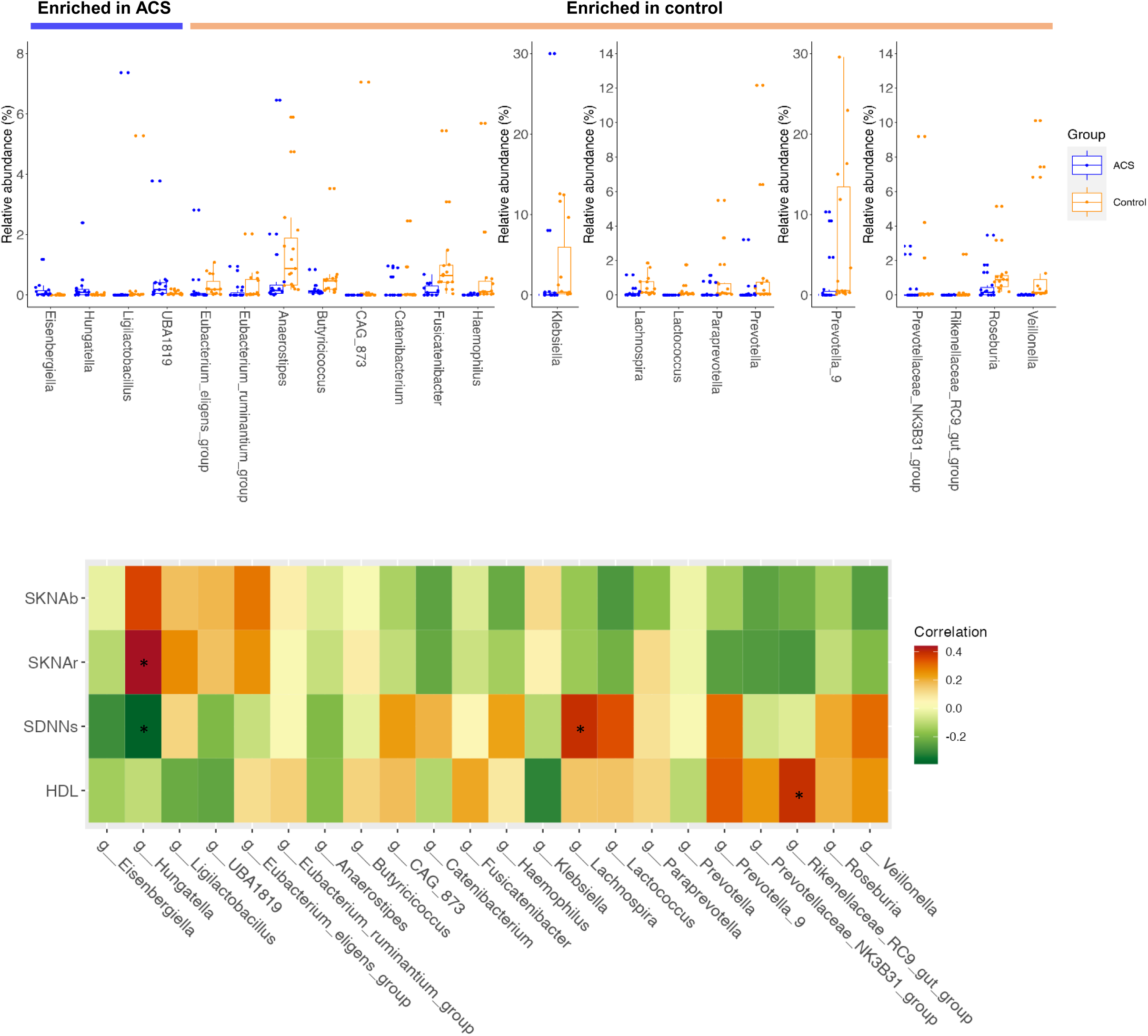
The 22 genera identified by intersection of LEfSe and random forest analysis. **A**, the relative abundances. **B**, the correlation with the SKNAb, SKNAr, SDNNs and HDL.

### Links between the gut microbiome and the clinical characteristics of ACS

Because HDL-C, SKNAb, SDNNs, and SKNAr demonstrated significant associations with the dissimilarity of gut microbiota between the ACS and control groups, we proceeded to explore the correlations among the microbial relative abundances of the 22 genera that were identified as discriminant taxa through the LEfSe and random forest analysis. As shown in Figure 4B, *Hungatella* exhibited a negative correlation with SDNNs (*r* = -0.392, *P* value = 0.032) and a positive correlation with SKNAr (*r* = 0.441, *P* value = 0.015). Notably, it also showed a marginally significant correlation with SKNAb (*r* = 0.358, *P* value = 0.052). *Lachnospira* was positively correlated with SDNNs (*r* = 0.393, *P* value = 0.032), while the Rikenellaceae RC9 gut group positively correlated with HDL-C (*r* = 0.392, *P* value = 0.032). None of the other discriminant taxa showed significant correlations with the four clinical parameters. Based on the results of LEfSe and random forest, we further analyzed the fixed effect to determine the correlation between SKNAb, SKNAr, and SDNNs and the abundance of gut microbiota after controlling for age and gender (Figure 5A and B). The fixed effects of SKNAb and SKNAr on *Hungatella* were 0.66 (*P* <0.001) and 5.757 (*P* <0.001), respectively. The fixed effect of SDNNs on *Lachnospira* was 0.057 (*P* <0.001).

**Figure 5.**
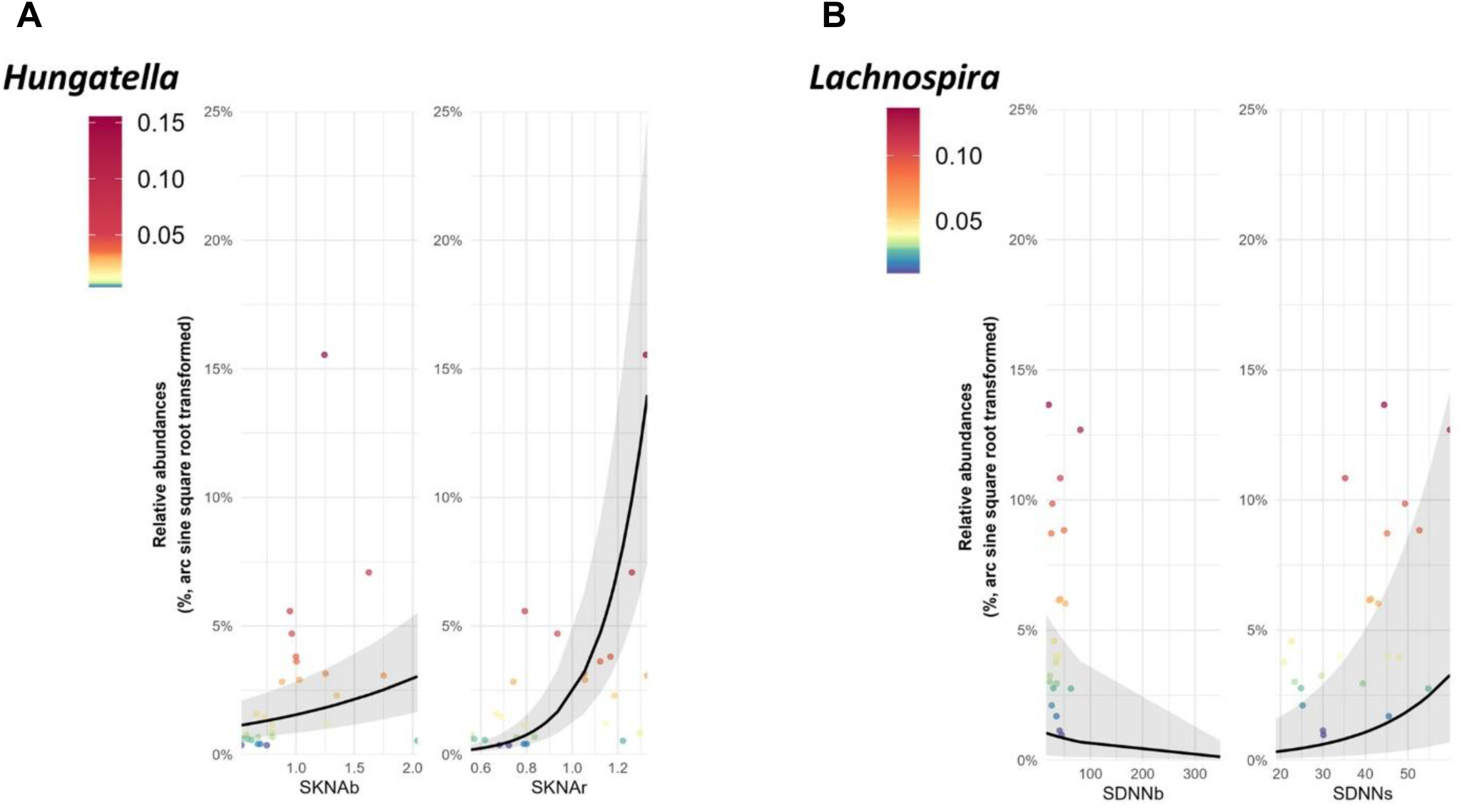
The fixed effect after controlling age and gender to determine the correlation between microbiota and autonomic survey. **A**, the correlation between *Hungatella* with SKNA. **B**, the correlation between *Lachnospira* with SDNN.

### Prediction of microbial functions

PICRUSt2 analysis showed that 49 and 41 MetaCyc pathways were enriched in the ACS and control groups, respectively (Figure 6). In the ACS group, 12 and 10 pathways were significantly enriched under amino acid biosynthesis and nucleoside and nucleotide biosynthesis. Among them, 7 pathways related to branched-chain amino acids (BCAA) biosynthesis were noted. Regarding the control group, 21 pathways were enriched under cofactor, prosthetic group, electron carrier, and vitamin biosynthesis, and 4 pathways under aromatic compound degradation.

**Figure 6.**
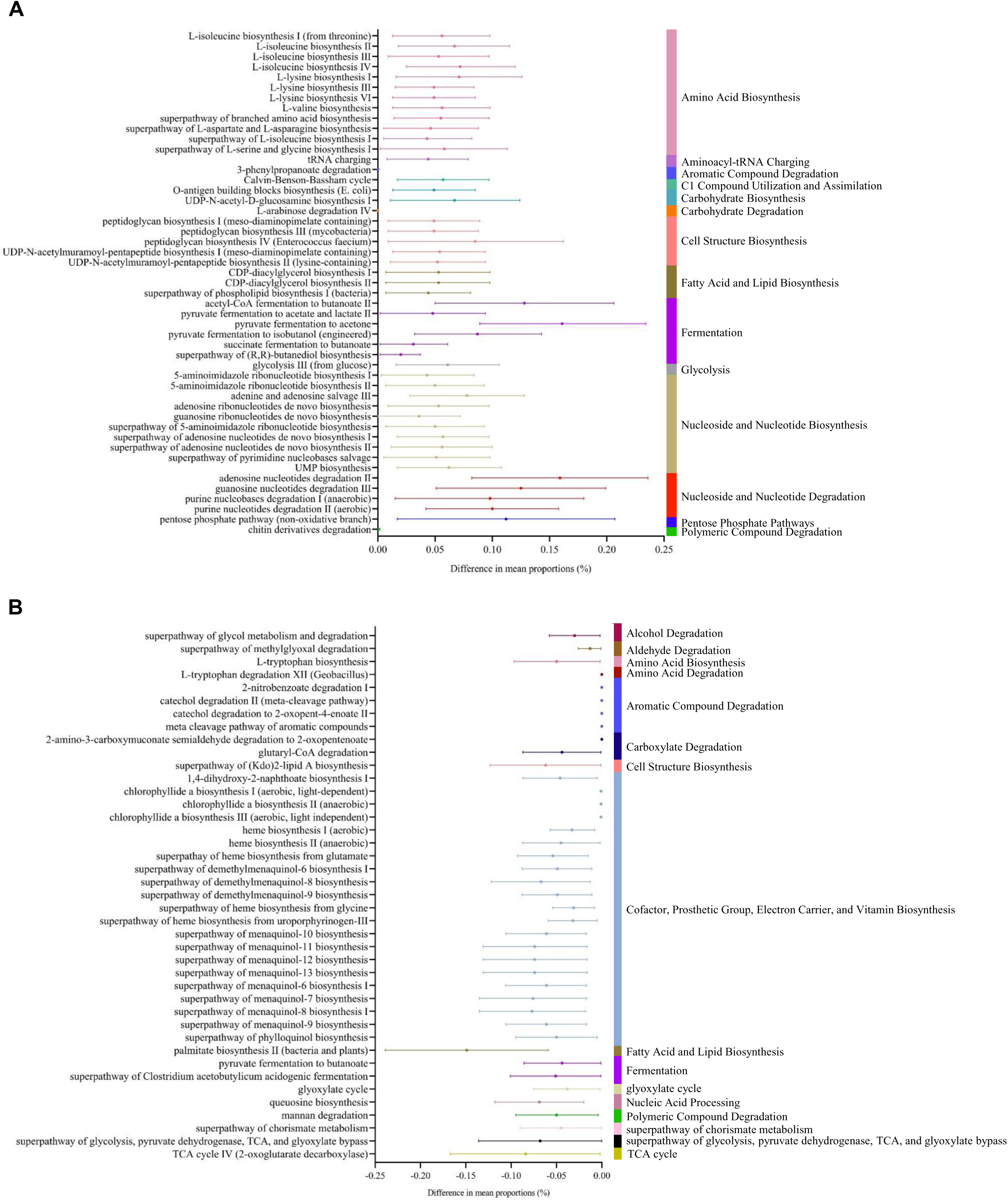
Comparison of Picrust2 predicted MetaCyc pathways between the ACS and control groups. **A**, The pathways enriched in the ACS group. **B**, The pathways enriched in the control group.

## DISCUSSION

While Meng et al. highlighted the important role of gut microbes-derived metabolite TMAO in the development of VA through the activation of cardiac SNS,^16^ our study is the first to demonstrate the interplay between skin SNS and gut dysbiosis in ACS patients. Previously, we have shown that SKNA was associated with VA in ACS, establishing it as a novel biomarker for ACS risk stratification.^12^ The current study further highlights the significant association of SKNA in ACS patients, specifically identifying a link with gut microbiota profiles.

The value of SKNA was positively correlated to the relative abundance of *Hungatella*, a TMA-producer (Figure 4 and Figure 5A). *Hungatella*, combined with *Eggerthella lenta*, was specifically required to convert L-carnitine to Trimethylamine (TMA).^32^ TMA is the diet-derived precursor to TMAO, the major metabolite of the microbiome correlated with the outcomes of ACS.^14^ Altered *Hungatella* abundance was linked to an increased incidence of unruptured intracranial aneurysms, an atherosclerotic cardiovascular disease.^33^ In two independent cross-sectional cohorts of patients with systolic heart failure (HF), increased *Hungatella* was associated with HF.^34^ *Hungatella* was also negatively correlated with HDL-C in stroke^35^ and positively correlated with IL-10 levels.^36^ Based on the above evidence, *Hungatella* is logically associated with cardiovascular disease and ACS. In addition, SGLT2i can reduce *Hungatella* in type 2 diabetes^37^ and polyphenol-rich extracts can lower plasma TMAO in overweight and obese adults in association with *Hungatella* modulation^38^, demonstrating the cardiovascular benefits of reducing *Hungatella* in subjects at risk of cardiovascular disease. Other than cardiovascular disease, *Hungatella* was also found to be more significant enrichment in patients with generalized anxiety disorder.^39^ As anxiety was thought to correlate with high SNS status, the association between *Hungatella* and SNS is reasonable.

Type 2 diabetes, often linked with gut dysbiosis, is prevalent among patients with ACS and is recognized as a risk factor for the development of ACS. However, Emoto et al. compared the composition of gut microbiota among coronary artery disease (CAD) patients, non-CAD controls but with coronary risk factors such as type 2 diabetes, and healthy volunteers to demonstrate that a change in gut microbiota in the CAD group was not associated with type 2 diabetes.^40^ A key aspect of our study was the exclusion of type 2 diabetes in the enrolled subjects to clearly define the changes in gut microbiota profiles specific to ACS. Compared to the matched controls, ACS patients exhibited a decrease in butyrate-producing bacteria and an increase in TMA-producing bacteria. These findings align with previous studies that included patients with type 2 diabetes,^13,15^ further highlighting the significant roles of these microbiota in the development of ACS.

Elevated production of BCAAs, including leucine, isoleucine, and valine, was observed in the predicted microbial functional analysis using PICRUSt2 (Figure 6). Among the 12 pathways related to amino acid biosynthesis that were elevated in ACS subjects, five were associated with isoleucine biosynthesis, one with valine biosynthesis, and one with overall BCAA biosynthesis. Previous studies have shown that increased plasma BCAA concentrations are found in patients with heart failure or coronary artery disease and can predict adverse outcomes.^17–19^ Beyond impaired cardiac BCAA catabolism, growing evidence suggested that abnormal BCAA accumulation may originate from the gut microbiota.^20^ Additionally, BCAA accumulation has been shown to exacerbate microglia-induced neuroinflammation in a rat model of ischemia/reperfusion.^41^ These findings further support the association between microbiota and SKNA in ACS, as addressed in this study.

This study has several limitations. First, the sample size was relatively small due to the rigorous selection process, particularly the exclusion of clinical factors such as type 2 diabetes and use of proton pump inhibitors or metformin, which are common in ACS patients. Although the stringent selection process gives the strength of getting rid of the influence of the clinical factor on microbiota data, it also reduce the external validity since the clinical factor is common in ACS patients. Another limitation was using the V3-V4 region of 16S rRNA gene sequencing, which limited the identification of some bacteria to the genus level only. Additionally, we did not measure TMAO or BCAA levels in the subjects, hindering us from evaluating their relationships with gut microbiota and SKNA. Finally, other factors beyond SKNAb, SKNAr, and SDNNs may have contributed to microbial diversity in our study, as indicated by the relatively low correlation coefficients (r = 0.151, 0.209, and 0.249, respectively, in Table 2).

In conclusion, this study is the first to demonstrate an association between SKNA and gut microbial profiles in ACS patients. By applying stringent selection criteria to exclude type 2 diabetes, we identified specific patterns in ACS patients, including alterations in butyrate-producing bacteria, TMA-producing bacteria, and gut microbial BCAA biosynthesis. Together with our previous research, these findings suggest that SKNA could serve as a potential biomarker for ACS.

## Nonstandard Abbreviations and Acronyms

ACS: Acute coronary syndrome
ANOSIM: Analysis of similarities
aSKNA: Average skin sympathetic nerve activity
ASVs: Amplicon sequence variants
BCAA: Branched-chain amino acids
CAD: Coronary artery disease
HRV: Heart rate variability
LAD: Linear discriminant analysis
LEfSe: Linear discriminant analysis effect size
PCoA: Principal coordinate analyses
PERMANOVA: Permutational multivariate analysis of variance
PICRUSt2: Phylogenetic investigation of communities by reconstruction of observed states 2
SDNN: Standard deviation of NN intervals
SDNNs: SDNN during stress phase
SG: Stellate ganglion
SKNA: Skin sympathetic nerve activity
SKNAb: SKNA parameters during baseline phase
SKNAr: SKNA parameters during recovery phase
SNS: Sympathetic nervous system
TMA: Trimethylamine
TMAO: Trimethylamine N-oxide
VA: Ventricular arrhythmias

## SOURCES OF FUNDING

This study was supported in part by grants from the Ministry of Science and Technology, R.O.C “MOST 110-2314-B-037-111” and ”MOST 113-2314-B-037 -054 -MY3”, Kaohsiung Medical University Hospital “KMUH111-1T10”, “SI11001”, “SI11101”, “SI11201”, “110KMUOR01”, “NK111P24”, “NSYSU-KMU-112-P13” and “KMUH111-1T10”.

## DISCLOSURES

None.

## Notes

### Competing Interest Statement

The authors have declared no competing interest.

## REFERENCES

1. Nascimento BR, Brant LCC, Marino BCA, Passaglia LG, Ribeiro ALP. Implementing myocardial infarction systems of care in low/middle-income countries. Heart. 2019;105:20–26. doi: 10.1136/heartjnl-2018-313398

2. Piccini JP, Schulte PJ, Pieper KS, Mehta RH, White HD, Van de Werf F, Ardissino D, Califf RM, Granger CB, Ohman EM, et al. Antiarrhythmic drug therapy for sustained ventricular arrhythmias complicating acute myocardial infarction. Critical care medicine. 2011;39:78–83. doi: 10.1097/CCM.0b013e3181fd6ad7

3. Shen MJ, Zipes DP. Role of the autonomic nervous system in modulating cardiac arrhythmias. Circulation research. 2014;114:1004–1021. doi: 10.1161/circresaha.113.302549

4. Chen PS, Chen LS, Fishbein MC, Lin SF, Nattel S. Role of the autonomic nervous system in atrial fibrillation: pathophysiology and therapy. Circulation research. 2014;114:1500–1515. doi: 10.1161/circresaha.114.303772

5. Malik M, Farrell T, Cripps T, Camm AJ. Heart rate variability in relation to prognosis after myocardial infarction: selection of optimal processing techniques. Eur Heart J. 1989;10:1060–1074. doi: 10.1093/oxfordjournals.eurheartj.a059428

6. Exner DV, Kavanagh KM, Slawnych MP, Mitchell LB, Ramadan D, Aggarwal SG, Noullett C, Van Schaik A, Mitchell RT, Shibata MA, et al. Noninvasive risk assessment early after a myocardial infarction the REFINE study. J Am Coll Cardiol. 2007;50:2275–2284. doi: 10.1016/j.jacc.2007.08.042

7. Kusayama T, Wong J, Liu X, He W, Doytchinova A, Robinson EA, Adams DE, Chen LS, Lin SF, Davoren K, et al. Simultaneous noninvasive recording of electrocardiogram and skin sympathetic nerve activity (neuECG). Nat Protoc. 2020;15:1853–1877. doi: 10.1038/s41596-020-0316-6

8. Kusayama T, Wan J, Doytchinova A, Wong J, Kabir RA, Mitscher G, Straka S, Shen C, Everett THt, Chen PS. Skin sympathetic nerve activity and the temporal clustering of cardiac arrhythmias. JCI Insight. 2019;4. doi: 10.1172/jci.insight.125853

9. Kabir RA, Doytchinova A, Liu X, Adams D, Straka S, Chen LS, Shen C, Lin SF, Everett THt, Chen PS. Crescendo Skin Sympathetic Nerve Activity and Ventricular Arrhythmia. Journal of the American College of Cardiology. 2017;70:3201–3202. doi: 10.1016/j.jacc.2017.10.065

10. Zhang P, Liang JJ, Cai C, Tian Y, Dai MY, Wong J, Everett THt, Wittwer ED, Barsness GW, Chen PS, et al. Characterization of skin sympathetic nerve activity in patients with cardiomyopathy and ventricular arrhythmia. Heart Rhythm. 2019;16:1669–1675. doi: 10.1016/j.hrthm.2019.06.008

11. Doytchinova A, Hassel JL, Yuan Y, Lin H, Yin D, Adams D, Straka S, Wright K, Smith K, Wagner D, et al. Simultaneous noninvasive recording of skin sympathetic nerve activity and electrocardiogram. Heart Rhythm. 2017;14:25–33. doi: 10.1016/j.hrthm.2016.09.019

12. Huang TC, Lin SJ, Chen CJ, Jhuo SJ, Chang CW, Lin SC, Chi NY, Chou LF, Tai LH, Liu YH, et al. Skin sympathetic nerve activity and ventricular arrhythmias in acute coronary syndrome. Heart Rhythm. 2022. doi: 10.1016/j.hrthm.2022.04.031

13. Amiri P, Hosseini SA, Ghaffari S, Tutunchi H, Mosharkesh E, Asghari S, Roshanravan N. Role of butyrate, a gut microbiota derived metabolite, in cardiovascular diseases: a comprehensive narrative review. Front Pharmacol. 2021;12:837509. doi: 10.3389/fphar.2021.837509837509837509 [pii]

14. Li XS, Obeid S, Klingenberg R, Gencer B, Mach F, Räber L, Windecker S, Rodondi N, Nanchen D, Muller O, et al. Gut microbiota-dependent trimethylamine N-oxide in acute coronary syndromes: a prognostic marker for incident cardiovascular events beyond traditional risk factors. Eur Heart J. 2017;38:814–824. doi: 10.1093/eurheartj/ehw582

15. Liu H, Chen X, Hu X, Niu H, Tian R, Wang H, Pang H, Jiang L, Qiu B, Chen X, et al. Alterations in the gut microbiome and metabolism with coronary artery disease severity. Microbiome. 2019;7:68. doi: 10.1186/s40168-019-0683-9

16. Meng G, Zhou X, Wang M, Zhou L, Wang Z, Wang M, Deng J, Wang Y, Zhou Z, Zhang Y, et al. Gut microbe-derived metabolite trimethylamine N-oxide activates the cardiac autonomic nervous system and facilitates ischemia-induced ventricular arrhythmia via two different pathways. EBioMedicine. 2019;44:656–664. doi: 10.1016/j.ebiom.2019.03.066

17. Grajeda-Iglesias C, Rom O, Aviram M. Branched-chain amino acids and atherosclerosis: friends or foes? Curr Opin Lipidol. 2018;29:166–169. doi: 10.1097/MOL.0000000000000494 00041433-201804000-00017 [pii]

18. White PJ, Newgard CB. Branched-chain amino acids in disease. Science. 2019;363:582–583. doi: 10.1126/science.aav0558363/6427/582 [pii]

19. McGarrah RW, White PJ. Branched-chain amino acids in cardiovascular disease. Nat Rev Cardiol. 2023;20:77–89. doi: 10.1038/s41569-022-00760-3 10.1038/s41569-022-00760-3 [pii]

20. Pedersen HK, Gudmundsdottir V, Nielsen HB, Hyotylainen T, Nielsen T, Jensen BA, Forslund K, Hildebrand F, Prifti E, Falony G, et al. Human gut microbes impact host serum metabolome and insulin sensitivity. Nature. 2016;535:376–381. doi: nature18646 [pii] 10.1038/nature18646

21. Kop WJ, Krantz DS, Nearing BD, Gottdiener JS, Quigley JF, O’Callahan M, DelNegro AA, Friehling TD, Karasik P, Suchday S, et al. Effects of acute mental stress and exercise on T-wave alternans in patients with implantable cardioverter defibrillators and controls. Circulation. 2004;109:1864–1869. doi: 10.1161/01.cir.0000124726.72615.60

22. Pan J, Tompkins WJ. A real-time QRS detection algorithm. IEEE transactions on bio-medical engineering. 1985;32:230–236. doi: 10.1109/tbme.1985.325532

23. Chan YH, Tsai WC, Shen C, Han S, Chen LS, Lin SF, Chen PS. Subcutaneous nerve activity is more accurate than heart rate variability in estimating cardiac sympathetic tone in ambulatory dogs with myocardial infarction. Heart Rhythm. 2015;12:1619–1627. doi: 10.1016/j.hrthm.2015.03.025

24. Hung WW, Peng P, Tsai YC, Jhou PS, Chang CC, Hsieh CC, Su YC, Dai CY, Hung WC. Gut microbiota compositions and metabolic functions in type 2 diabetes differ with glycemic durability to metformin monotherapy. Diabetes Res Clin Pract. 2021;174:108731. doi: S0168-8227(21)00084-X [pii] 10.1016/j.diabres.2021.108731

25. Bolyen E, Rideout JR, Dillon MR, Bokulich NA, Abnet CC, Al-Ghalith GA, Alexander H, Alm EJ, Arumugam M, Asnicar F, et al. Reproducible, interactive, scalable and extensible microbiome data science using QIIME 2. Nat Biotechnol. 2019;37:852–857. doi: 10.1038/s41587-019-0209-9 10.1038/s41587-019-0209-9 [pii]

26. Callahan BJ, McMurdie PJ, Rosen MJ, Han AW, Johnson AJ, Holmes SP. DADA2: High-resolution sample inference from Illumina amplicon data. Nat Methods. 2016;13:581–583. doi: 10.1038/nmeth.3869nmeth.3869 [pii]

27. Xia Y, Sun J. Hypothesis testing and statistical analysis of microbiome. Genes Dis. 2017;4:138–148. doi: 10.1016/j.gendis.2017.06.001 S2352-3042(17)30035-1 [pii]

28. Pedregosa F, Varoquaux G, Gramfort A, Michel V, Thirion B, Grisel O, Blondel M, Prettenhofer P, Weiss R, Dubourg V. Scikit-learn: Machine learning in Python. the Journal of machine Learning research. 2011;12:2825–2830.

29. Quast C, Pruesse E, Yilmaz P, Gerken J, Schweer T, Yarza P, Peplies J, Glöckner FO. The SILVA ribosomal RNA gene database project: improved data processing and web-based tools. Nucleic Acids Research. 2012;41:D590–D596. doi: 10.1093/nar/gks1219

30. Segata N, Izard J, Waldron L, Gevers D, Miropolsky L, Garrett WS, Huttenhower C. Metagenomic biomarker discovery and explanation. Genome Biology. 2011;12:R60. doi: 10.1186/gb-2011-12-6-r60

31. Douglas GM, Maffei VJ, Zaneveld JR, Yurgel SN, Brown JR, Taylor CM, Huttenhower C, Langille MGI. PICRUSt2 for prediction of metagenome functions. Nature Biotechnology. 2020;38:685–688. doi: 10.1038/s41587-020-0548-6

32. Koeth RA, Lam-Galvez BR, Kirsop J, Wang Z, Levison BS, Gu X, Copeland MF, Bartlett D, Cody DB, Dai HJ, et al. l-Carnitine in omnivorous diets induces an atherogenic gut microbial pathway in humans. The Journal of clinical investigation. 2019;129:373–387. doi: 10.1172/jci94601

33. Li H, Xu H, Li Y, Jiang Y, Hu Y, Liu T, Tian X, Zhao X, Zhu Y, Wang S, et al. Alterations of gut microbiota contribute to the progression of unruptured intracranial aneurysms. Nature communications. 2020;11:3218. doi: 10.1038/s41467-020-16990-3

34. Kummen M, Mayerhofer CCK, Vestad B, Broch K, Awoyemi A, Storm-Larsen C, Ueland T, Yndestad A, Hov JR, Trøseid M. Gut Microbiota Signature in Heart Failure Defined From Profiling of 2 Independent Cohorts. Journal of the American College of Cardiology. 2018;71:1184–1186. doi: 10.1016/j.jacc.2017.12.057

35. Cui W, Xu L, Huang L, Tian Y, Yang Y, Li Y, Yu Q. Changes of gut microbiota in patients at different phases of stroke. CNS Neuroscience & Therapeutics. 2023;29:3416–3429. doi: 10.1111/cns.14271

36. Sun H, Gu M, Li Z, Chen X, Zhou J. Gut Microbiota Dysbiosis in Acute Ischemic Stroke Associated With 3-Month Unfavorable Outcome. Frontiers in Neurology. 2021;12:799222. doi: 10.3389/fneur.2021.799222

37. Deng X, Zhang C, Wang P, Wei W, Shi X, Wang P, Yang J, Wang L, Tang S, Fang Y, et al. Cardiovascular Benefits of Empagliflozin Are Associated With Gut Microbiota and Plasma Metabolites in Type 2 Diabetes. J Clin Endocrinol Metab. 2022;107:1888–1896. doi: 10.1210/clinem/dgac210

38. Rehman A, Tyree SM, Fehlbaum S, DunnGalvin G, Panagos CG, Guy B, Patel S, Dinan TG, Duttaroy AK, Duss R, et al. A water-soluble tomato extract rich in secondary plant metabolites lowers trimethylamine-n-oxide and modulates gut microbiota: a randomized, double-blind, placebo-controlled cross-over study in overweight and obese adults. Journal of Nutrition. 2023;153:96–105. doi: 10.1016/j.tjnut.2022.11.009

39. Chen YH, Bai J, Wu D, Yu SF, Qiang XL, Bai H, Wang HN, Peng ZW. Association between fecal microbiota and generalized anxiety disorder: Severity and early treatment response. Journal of Affective Disorders. 2019;259:56–66. doi: 10.1016/j.jad.2019.08.014

40. Emoto T, Yamashita T, Sasaki N, Hirota Y, Hayashi T, So A, Kasahara K, Yodoi K, Matsumoto T, Mizoguchi T, et al. Analysis of gut microbiota in coronary artery disease patients: a possible link between gut microbiota and coronary artery disease. J Atheroscler Thromb. 2016;23:908–921. doi: 10.5551/jat.32672

41. Shen J, Guo H, Liu S, Jin W, Zhang ZW, Zhang Y, Liu K, Mao S, Zhou Z, Xie L, et al. Aberrant branched-chain amino acid accumulation along the microbiota-gut-brain axis: Crucial targets affecting the occurrence and treatment of ischaemic stroke. Br J Pharmacol. 2023;180:347–368. doi: 10.1111/bph.15965

